# Neural response variability and divisive normalization

**DOI:** 10.1101/508838

**Authors:** Ruben Coen-Cagli, Selina S. Solomon

## Abstract

Cortical responses to repeated presentations of a stimulus are variable. This variability is sensitive to experimental manipulations that are also known to engage divisive normalization: a widespread description of neural activity as the ratio of a numerator (the excitatory stimulus drive) and denominator (the normalization signal). Yet, we lack a framework to quantify the effects of normalization on response variability. We extended the standard normalization model, treating the numerator and denominator as stochastic quantities, and derived a method to infer the single-trial normalization strength, which cannot be measured directly. The model revealed a general reduction of response variability in macaque primary visual cortex for neurons that were more strongly normalized, and during trials in which normalization was inferred to be strong. This framework could enable a direct quantification of the impact of single-trial normalization on perceptual judgments, and can readily be applied to other sensory and non-sensory factors.

## Introduction

Variability is a widespread feature of neuronal activity in sensory cortex: spike counts in response to repeated presentations of the same sensory input are variable (Arieli et al. 1996; Tolhurst et al. 1983). Traditionally, studies of neural coding have focused on the mean response averaged over repetitions, i.e. the firing rate, to extract the signal (differences in activity between conditions) from the noise (changes across repeats of the same condition). In this view variability is a disturbance of neural coding. However, just like firing rate, variations of neural activity across trials also contain meaningful information. For instance, stimulus onset reduces variability, compared to spontaneous activity, in many cortical areas (Churchland et al., 2010). In early visual cortex, variability is modulated along several stimulus dimensions, e.g. contrast (Orbán et al. 2016), size (Snyder et al. 2014), orientation (Goris et al. 2014), and motion direction (Ponce-Alvarez et al. 2013). Non-sensory signals like attention (Cohen & Maunsell, 2009; Ecker et al., 2014; Harris & Thiele, 2011; Mitchell et al. 2009; Rabinowitz et al. 2015) and locomotion (Bennett et al. 2013; Dadarlat & Stryker, 2017) also modulate neuronal variability. Many of these effects persist even after accounting for firing rate differences (Churchland et al., 2010), suggesting variability may be a genuine coding dimension of neuronal activity beyond firing rate.

Therefore, it is important to understand how rate and variability are jointly modulated, but unfortunately, existing models have a limited ability to do so. This is because successful descriptive models of neuronal variability (Charles et al. 2018; Goris et al., 2014; Paninski, 2004; Pillow et al., 2008) typically oversimplify the nonlinear stimulus dependence of cortical responses. An accurate description of such nonlinear behavior includes divisive normalization (Albrecht & Geisler, 1991; Heeger, 1992), a canonical operation evident across multiple brain areas (Carandini & Heeger, 2011): the firing rate of a neuron is the ratio between the stimulus drive to its receptive field (the numerator), and the stimulus drive to an ensemble of other neurons (the denominator, termed normalization signal). However, typically normalization models describe the firing rate, and add noise (often implicitly, i.e. for data fitting) only to the output of the normalization equation, so the variability has no dependence on the normalization: therefore, they cannot address whether and how normalization impacts variability.

On the other hand, there is mounting evidence that normalization may have an impact on variability. First, factors that are found experimentally to affect response variability, such as stimulus contrast (Orbán et al., 2016), stimulus size (Snyder et al., 2014) and attentional modulation (Cohen & Maunsell, 2009; Mitchell et al. 2009; Rabinowitz et al., 2015), are also thought to control the strength of normalization (Coen-Cagli et al. 2015; Heeger, 1992; Reynolds & Heeger, 2009; Schwartz & Simoncelli, 2001). Second, recent work has used numerical simulations to demonstrate that normalization can strongly modulate one aspect of response variability, namely how this variability is shared between pairs of neurons (i.e. noise correlations) (Tripp, 2012; Verhoef & Maunsell, 2017). Furthermore, although the mechanisms of normalization are not fully known, a successful network model has shown that inhibitory stabilization of network activity (Hennequin et al. 2018; Rubin et al. 2015) recapitulates many effects typically described by normalization, further pointing to a link between changes in effective normalization strength and changes in variability, via network stabilization.

To better understand and quantify the relation between normalization and variability, it is necessary to have a descriptive model that explicitly parametrizes this relation, and allows one to estimate those parameters from data. Furthermore, to investigate how normalization might affect perception through its effects on variability, it is necessary to compare individual perceptual choices to the instantaneous strength of normalization, which unfortunately cannot be measured directly.

To achieve these goals, we first derived new mathematical results linking normalization and the across-trial spike count distribution. We treated the numerator and the normalization signal as two random variables, and derived analytical estimators for the across-trial mean and variance of their ratio, as well as an efficient method to infer the single-trial value of the normalization signal. The model, termed ratio of Gaussians (RoG), reproduced Poisson-like behavior and rate-dependent Fano factors consistent with known cortical data (Goris et al., 2014; Tolhurst et al., 1983). We then applied the RoG model to contrast-tuning data recorded in macaque primary visual cortex (V1), because contrast modulation of firing rate is well described by the standard normalization model (Heeger, 1992; Tolhurst & Heeger, 1997). Our model captured accurately both the mean and variance of the spike count as a function of contrast, including a systematic reduction of the Fano factor at high contrast that has been reported recently (Orbán et al., 2016) and that eluded other popular descriptive models. Furthermore, the RoG predicted that increasing the strength of the normalization signal has a general stabilizing effect on neuronal responses. We tested this prediction, and found that V1 activity is more stable for neurons that are more strongly normalized, as well as during single trials in which the normalization signal is inferred to be stronger.

In summary, we have developed the RoG model and validated it on V1 contrast response data. The model highlights a general relation between divisive normalization and response variability, and can be extended to quantify similar relations for other stimulus dimensions and non-sensory factors, beyond stimulus contrast in V1, as well as to study the impact of single-trial normalization strength on perception.

## Results

We developed a descriptive model to quantify how normalization affects response variability. To this aim, we extended the standard normalization model, in which the spike count of a neuron is described by the ratio between the stimulus drive to the neuron (the numerator, *N*) and the normalization signal (the denominator, *D*). We treated both *N* and *D* as random variables (Fig. 1a). Specifically, we assumed Gaussian noise in both *N* and *D*, and thus obtained a ratio of two Gaussian variables (RoG). Note that in this model the mean and variance of the spike count are not independent: they both follow from the statistics of *N* and *D*. There is no simple closed-form expression for the RoG distribution, but if *D* has negligible probability mass at or below 0 the RoG is closely approximated by a Gaussian (Marsaglia, 1965, 2006; Pham-Gia et al. 2006). Under this condition, we derived analytical expressions to approximate the mean and variance of the RoG (Methods eq. (1.4); Fig. 1b).

**Figure 1.**
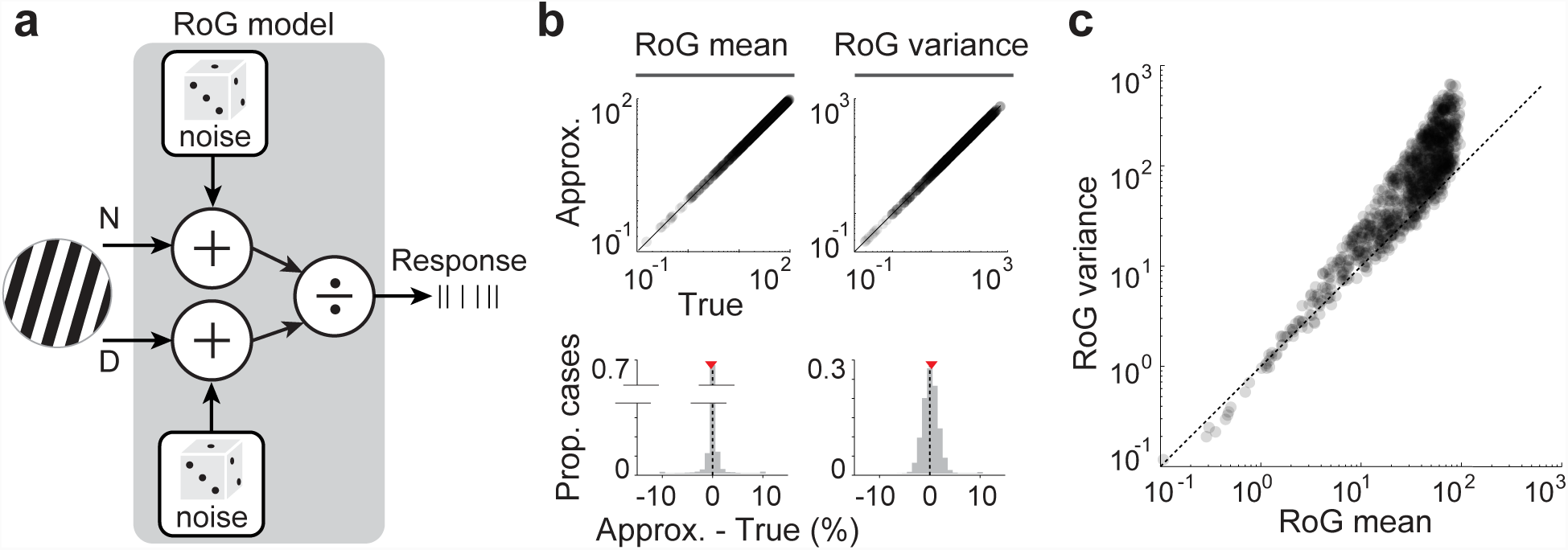
Ratio of Gaussians model. (**a**) The RoG model describes neural responses as the ratio of two stimulus-driven signals: the numerator *N* and denominator, or normalization signal, *D*. Both *N* and *D* are corrupted by additive Gaussian noise with Poisson-like variability (Methods eq. (1.9)). (**b**) Comparison of the approximation we derived for the RoG mean and variance (Methods eq. (1.4)), and the true value (estimated from 1,000,000 simulated trials), over 1,000 experiments. Each experiment uses a different set of model parameters (i.e. corresponding to different neurons and stimuli), and each trial is a random draw from the corresponding distribution. Red triangles: average percent difference between true and approximated mean (left) or variance (right). (**c**) RoG variance versus mean across 1,000 simulated experiments. Model parameters for the simulations in (**b**) and (**c**) are drawn uniformly in the following intervals:

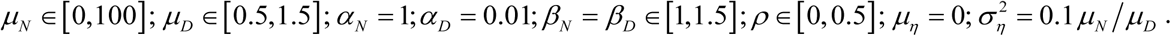

### The RoG model can capture the typical relation between response mean and variance

Cortical responses are characterized by a monotonic relation between the variance and mean of the spike count, known as Poisson-like variability (Goris et al., 2014; Tolhurst et al., 1983). Most existing models include this relation explicitly (Charles et al., 2018; Ecker et al., 2014; Goris et al., 2014; Lin et al. 2015; Paninski, 2004; Pillow et al., 2008). Differently from those models, the RoG encompasses a broader range of mean-variance relations, therefore we first asked if the RoG model too can capture Poisson-like variability. To constrain the RoG, we reasoned that if *N* and *D* describe signals of neural origin, each one of them should be Poisson-like (in particular, *D* is usually thought of as the summed activity of many neurons, so also Poisson-like). Therefore, we assumed a power-law relation between the mean and variance (Carandini, Heeger, & Movshon, 1997; Tolhurst et al., 1983) for both *N* and *D*. With these choices, the RoG model also displayed Poisson-like variability across a broad range of values for the power-law parameters, and the parameters could be easily tuned to obtain variance-to-mean relations qualitatively consistent with those typically observed in V1 (Fig. 1c).

Therefore, the RoG model can be easily constrained to capture Poisson-like variability and has the advantage, over other models, that it models explicitly the divisive normalization form of cortical firing rates.

### Contrast modulation of V1 variability is best described by the RoG model

To test the RoG model quantitatively, we recorded responses of macaque V1 neurons (n=172) to sinusoidal gratings drifting in 4 directions at 5 contrast levels between 6.25% and 100%. Each grating was presented for 500-800ms and repeated 20-25 times. As our interest is on the relationship between normalization and variability, for each neuron we analyzed only the responses to the best direction out of those presented and we focused instead on contrast manipulation because normalization is known to provide excellent fits to the contrast response function (Heeger, 1992; Tolhurst & Heeger, 1997). We therefore parametrized the means of *N* and *D* in our model as in the standard normalization model (eq. (1.16)). Namely, we assumed *N* proportional to the squared contrast, and *D* equal to the squared contrast plus a constant. We then optimized jointly the parameters for the means of *N* and *D*, as well as those for the power-law relation between the means and variances, by maximum likelihood. Figure 2a,b shows for two example neurons that the RoG model captured accurately both the mean count (as expected) as well as the dependence of the variance on contrast. We quantified the predictive power of the model by the cross-validated goodness of fit (details in Methods), i.e. the log-likelihood of a test dataset not used for model training, normalized between a null model (goodness of fit = 0) and the oracle model (goodness of fit = 1). Across all neurons, the median cross-validated goodness of fit was 0.85 (c.i. [0.80 0.87]), denoting excellent predictive power.

**Figure 2.**
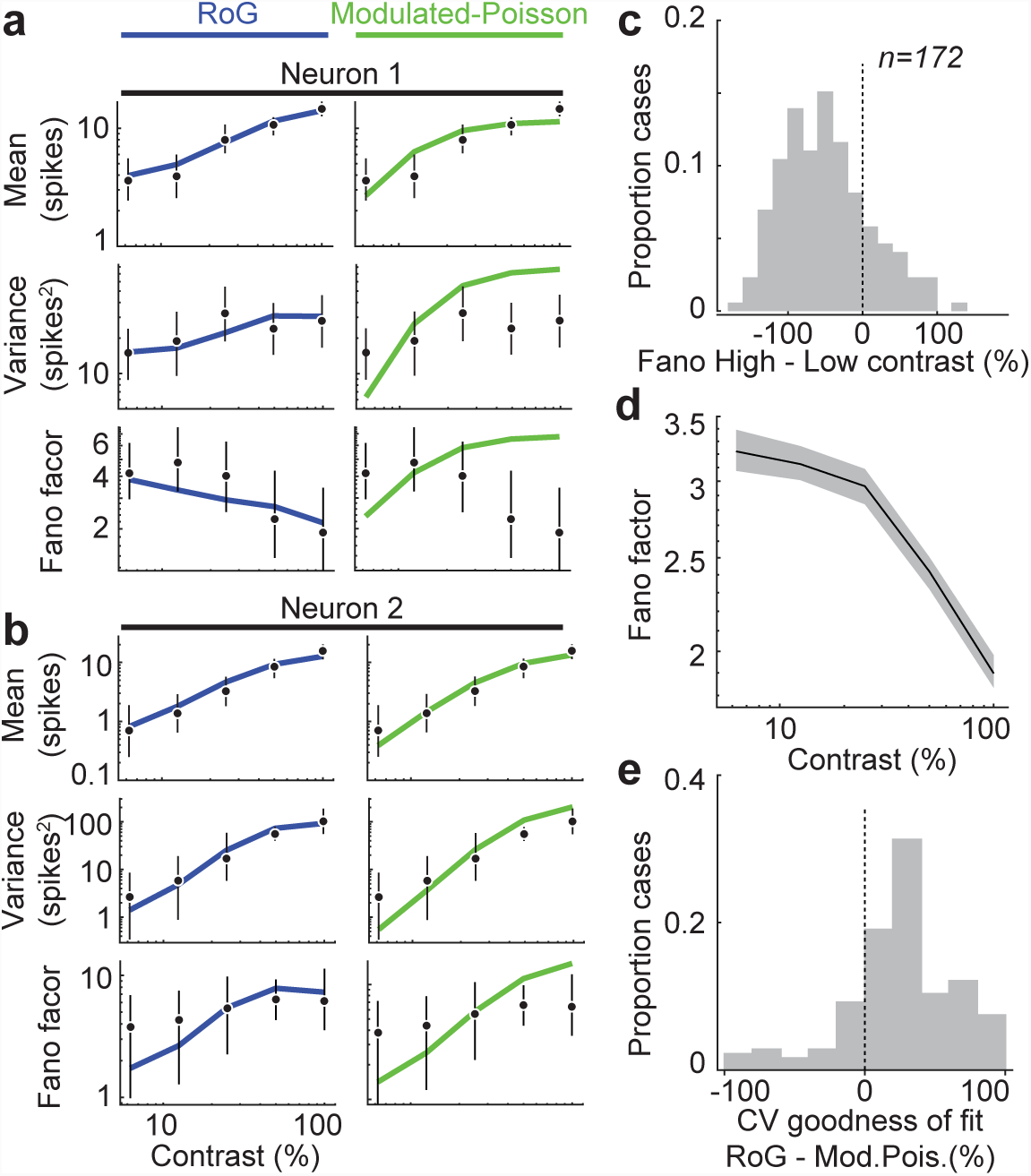
RoG model captures contrast modulation of firing rate and variability. (**a**,**b**) Mean spike count (top), variance (middle), and Fano factor (bottom) as a function of contrast for two example neurons. Circles and black lines: data and 95% c.i. Blue lines: RoG model fits. Green lines: Modulated-Poisson model fits. (**c**) Histogram of percent difference between Fano factors at high versus low contrast. (**d**) Geometric mean (black line) and 95% c.i. (gray shaded area) of the Fano factor across neurons, as a function of contrast. (**e**) Percent difference in cross-validated goodness of fit between the RoG and the Modulated-Poisson models. See Methods for details on model fitting and evaluation.

We observed a marked decrease in Fano factor at higher contrasts in the example neuron of Fig. 2a, also consistent with a previous report (Orbán et al., 2016). This was because the variance increased with contrast more slowly than the mean rate (Fig. 2a). Across neurons, high contrast stimuli elicited lower Fano factors compared to low contrast stimuli for 138/172 neurons (average Fano at low contrast 3.2, c.i. [2.9 3.5]; at high contrast 1.9, c.i. [1.7 2.0]), and the average Fano decreased monotonically with contrast (Fig. 2c,d). This behavior was captured by the RoG model (the blue curves in Fig. 2a), although we note that the model could also capture the opposite effect in the small subset of neurons (n=34/172) that displayed it (e.g. Fig. 2b). The RoG can capture both increases and decreases of Fano factor with contrast because contrast affects the means of both *N* and *D*, which can have opposing effects on Fano (eq. (1.8)). In addition, the subset of neurons whose Fano factor increased with contrast had lower spontaneous Fano factors and maximum firing rate, and larger best-fit exponents for the power law relations of *N* and *D* (the other parameters did not change substantially; Table 1).

**Table 1.** Comparison of neurons with Fano factor increasing versus decreasing with contrast. P-values are for the difference between the means of the two subpopulations. For Fano factors, p-values are computed on the logarithm. Columns with bold numbers have significant difference. Because these analyses use the best-fit parameters of the RoG model, they were performed on the subsets of neurons with goodness of fit > 0.5 (32/34 neurons for the first row; 120/138 neurons for the second row).

| | Full-contrast<br>Fano | Spontaneous<br>Fano | $R_{\max}$ | $\epsilon$ | $\alpha_N$ | $\alpha_D$ | $\beta_N$ | $\beta_D$ |
| --- | --- | --- | --- | --- | --- | --- | --- | --- |
| Fano increasing with<br>contrast (n=34) | 2.40 | **1.55** | **5.8** | 54.8 | 11.3 | 12.6 | **1.70** | **1.60** |
| Fano decreasing with<br>contrast (n=138) | 1.87 | **3.56** | **10.9** | 38.7 | 8.7 | 10.1 | **1.53** | **1.30** |
| p-value (different<br>means) | 0.04 | **$2 \cdot 10^{-10}$** | **0.0003** | 0.02 | 0.13 | 0.11 | **0.003** | **$9 \cdot 10^{-6}$** |

The main effect of contrast on Fano factors was qualitatively at odds with the Poisson model (constant Fano of 1), and also with a popular extension, the modulated-Poisson model (Goris et al., 2014). In the modulated-Poisson model, the variance increases faster than the mean and therefore the Fano factor could only increase with contrast, contrary to what we observed in the data (Fig. 2c,d). To quantify this discrepancy, we fitted an alternative model in which the mean was parametrized as in the RoG model, but the variability was modulated-Poisson (see Methods, Alternative Models). As expected, the alternative model could not describe well the variance data, particularly when the Fano factor decreased with contrast (e.g. green curves in Fig. 2a), and the fit quality (median cross-validated goodness of fit 0.62, c.i. [0.59 0.63]) was worse than the RoG model for most neurons (Fig. 2e). The RoG achieved a higher fit quality (0.79 vs. 0.66, p=0.023) also for the small subset of neurons whose Fano factor increased with contrast (e.g. Fig. 2b). We note that the modulated-Poisson model may be easily rescued with additional free parameters, i.e. assuming that the variance of the gain modulator is itself contrast-dependent and is reduced by high-contrast stimulation (Goris et al. 2017).

These results indicate that the RoG model successfully extends divisive normalization, to capture simultaneously the stimulus dependence of spike count mean and variance. Therefore, this model allows us to study directly how normalization affects variability.

### Normalization often reduces response variability

We have shown above that increasing stimulus contrast often decreases Fano factors. Moreover, in the standard normalization formulation, increasing stimulus contrast also increases the strength of the normalization signal. We then asked whether this inverse relation between strength of normalization and Fano factor is general, i.e. whether increasing normalization always contributes to stabilizing the output of the RoG model. The explicit normalization form of the RoG model allowed us to answer this question by studying directly the relation between normalization and variability.

We first analyzed, in the model, the dependence of the RoG variance on the mean of *D*, when everything else is held constant (including the variance of *D*). We found analytically that, if *N* and *D* are uncorrelated, the RoG variance can only decrease when the mean of *D* increases (Methods eq. (1.4)). Increasing the mean of *D* also decreases the RoG mean, therefore in principle an increase in *D* could have different effects on the Fano factor. However, we found analytically that the RoG mean always decreased slower than the variance, and so increasing the mean of *D* always reduced the Fano factor (Methods eq. (1.6)). We further verified that these results hold also if *N* and *D* are correlated, except for extremely large correlation values (Methods eq. (1.7)). We thus conclude that increasing normalization strength, when everything else is constant, usually stabilizes the RoG output.

We then considered the case with power-law relation between the mean and variance for *N* and *D*. In this case, increasing the mean of *D* increases its variance, and so the effects on the RoG variance and Fano need not be the same as in the previous analysis. Because no simple analytical solutions could be found for this case, we ran simulations in which we varied the mean of *D* systematically while keeping the mean of *N* constant, and tested different values of the parameters of the variance-to-mean relation for *N* and *D*. We observed a strong trend for reduced Fano factors as the mean of *D* increased, across a broad range of parameter settings (Fig. 3a), although this trend could be reversed for larger values of the exponent in the power-law of *D*, or if the additive noise has a relatively large Fano factor (see eqs. (1.8),(1.9) and related text).

**Figure 3.**
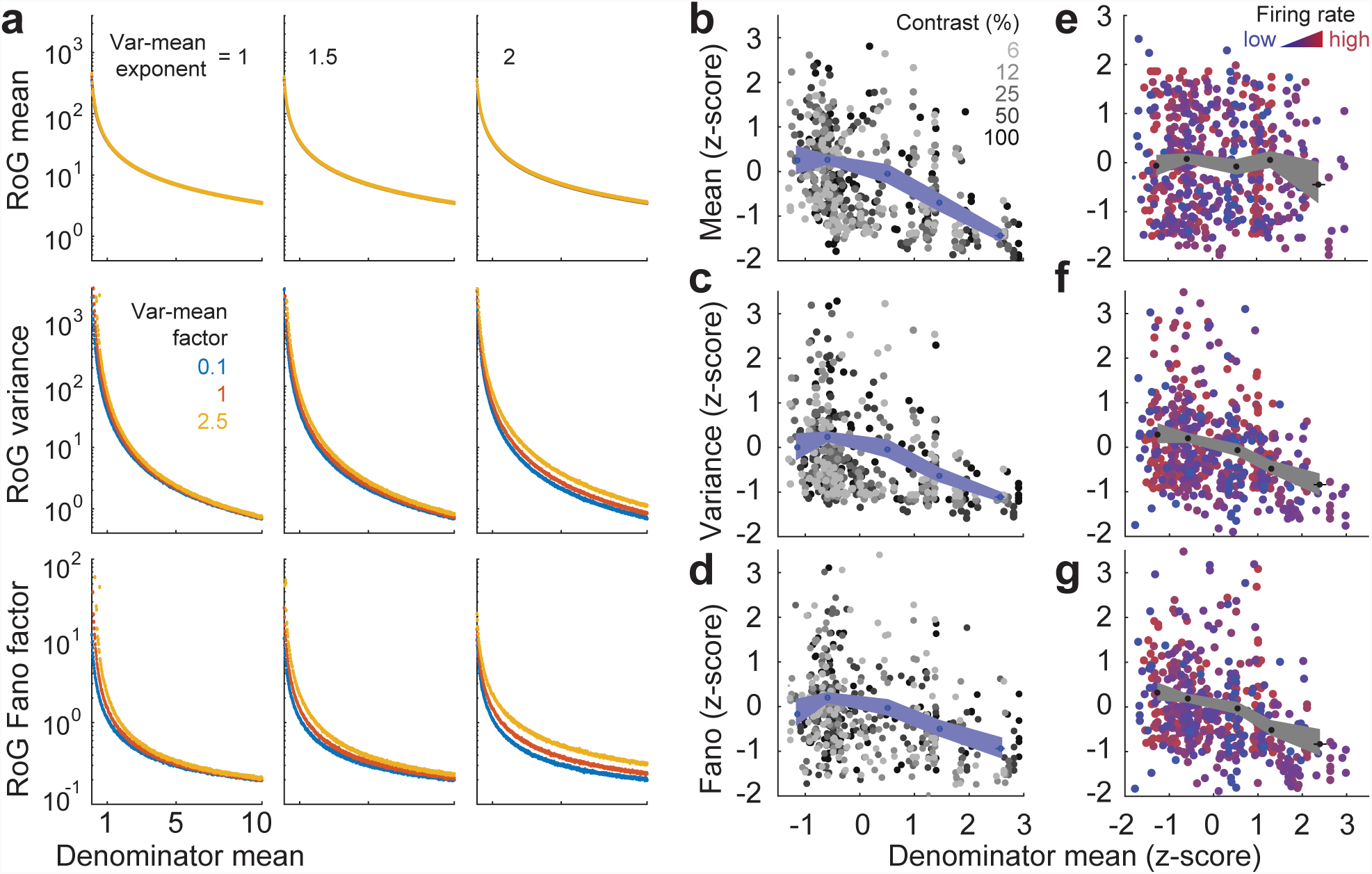
Normalization reduces response variability. (**a**) Relation between the normalization strength (abscissa) and mean (ordinate, top), variance (middle), or Fano factor (bottom) in the RoG model. Columns: different values of the exponent of the relation between mean and variance of the denominator. Different colors denote different values of the proportionality factor between mean and variance of the denominator. Other parameters are set to: 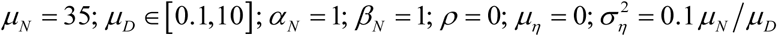. (**b-d**) Relation between the z-scored normalization strength and the z-scored response mean (**b**), variance (**c**), or Fano factor (**d**) in the data. Z-scoring was performed across all neurons, but separately within each contrast condition. Gray symbols denote individual neurons and contrasts. Blue symbols denote average across neurons and contrasts; shaded areas: 95% c.i. (**e-g**) Similar to (**b-d**), but with z-scoring performed across all neurons and contrast levels with similar firing rate. Firing rate bins were logarithmically spaced between the minimum and maximum firing rates measured. Black symbols denote average across neurons and contrasts; shaded areas: 95% c.i.

This analysis thus suggests that, in the RoG model, normalization often reduces spike count variability and Fano factors. If this is the case also in V1, we would expect that in our dataset neurons that are more strongly normalized are also less variable. To test the prediction, we first computed for each neuron and each contrast level the mean of *D*, i.e. the normalization strength, from the best fit parameters. Because the analysis relies on the fit parameters, we only included the 152/172 neurons with goodness-of-fit > 0.5. Different from the model analysis above, in which we varied *D* while keeping *N* constant, in the V1 data both *D* and *N* varied across neurons and contrast levels. Therefore, to isolate the effect of *D*, we performed two complementary analyses. First, we analyzed each contrast level separately and we z-scored the strength of *D* across neurons. We found that neurons with stronger normalization tended to have lower mean as expected (Fig. 3b) and even lower variance (Fig. 3c), resulting often in lower Fano factors (Fig. 3d). Across neurons and contrast levels, there was a significant negative correlation between the z-score of Fano and of *D* (Spearman correlation = −0.29, p=6*10^−16^). The negative correlation was significant also at each individual contrast level (p<0.03) except at 50% contrast (p=0.68). Second, we used mean-matching (Churchland et al., 2010) to assess the relation between normalization strength and variability, independent of firing rate. Specifically, we binned, across neurons and contrast levels, all cases with similar firing rate and then computed z-scores within each bin. In this case, as expected, there was no relation between the firing rate and *D* (Fig. 3e; Spearman correlation = −0.02, p=0.6), but we observed a strong negative correlation between the variance and *D* (Fig. 3f; Spearman correlation = −0.32, p=3*10^−19^) and between the Fano and *D* (Fig. 3g; Spearman correlation = −0.38, p=9*10^−27^).

Therefore, the RoG model predicts a general, inverse relation between the average normalization strength and the variability of cortical responses, which is apparent in V1 activity.

### Inference of single-trial normalization strength

The RoG model extends the classical normalization model, allowing it to simultaneously capture the mean and variance of the spike count across trials. An important advantage of the RoG formulation is that, because it treats the normalization signal explicitly as a random variable, it allows for inference of the normalization strength in single-trial activity—a quantity that cannot be measured directly with current experimental techniques. Therefore, we derived analytical expressions for the inferred value of the single-trial normalization strength (i.e. the most probable value of *D* for the observed spike count, according to the RoG model) and its uncertainty. Intuitively, if we knew the single-trial value of the numerator *N*, then we could estimate *D* simply as the ratio between *N* and the measured spike count. However, because *N* is not measurable either, we have to compute a so-called posterior probability distribution of the possible values of *D*, which amounts roughly to combining different estimates of *D* corresponding to all possible values of *N* (termed marginalization). This posterior distribution is computed via a direct application of Bayes rule, i.e. combining the evidence (the likelihood of the observed spike count for different possible values of *D*, under the RoG model) with the prior (i.e. the probability of the expected values of *D*, before observing the spike count; Methods eq. (1.10)). Note that the prior distribution of *D* is, by the definition of the RoG model, a Gaussian per eq. (1.2) whose (prior) mean and variance are estimated separately by model fitting across trials as shown above. The posterior distribution turned out to be Gaussian also, which allowed us to derive simple closed-form expressions for its (posterior) mean and variance (Methods eq. (1.12)), corresponding respectively the inferred value of *D* and the associated uncertainty.

We first validated these estimators in 10,000 simulated experiments, each with a different set of parameters for *N* and *D*. The example in Fig. 4a shows one experiment in which, across 100 simulated trials, estimates of *D* were strongly correlated with the ground-truth value. Across these simulated experiments however we observed a broad range of correlations (Fig. 4b,c), including cases with non-significant correlation. We reasoned that, if the prior variance of *D* is much smaller than that of *N*, the inferred single-trial value of *D* cannot be accurate. Intuitively, this is because when the prior variance of *D* is small, the single-trial value of *D* is always very close to the mean, i.e. it is almost constant across trials, therefore across-trial changes in the ratio are largely determined by across-trial changes in *N*. Therefore, while the average of *D* can be estimated accurately, its single-trial variations around the average cannot. This is also evident from eq. (1.12), where in the limit that the variance of *N* is large, the single-trial estimate of *D* becomes equal to its prior mean and independent of the measured spike count in that trial. Conversely, when the variance of *D* is larger relative to *N*, then the inference of *D* should be more accurate. In agreement with this, we found in our simulations that the quality of the inferred values of *D* increased systematically with the ratio between the variances of *D* and *N* (Fig. 4d). We note however that the estimates remained unbiased on average (within 0.01% across all simulated experiments) regardless of the ratio of variances (Fig. 4e).

**Figure 4.**
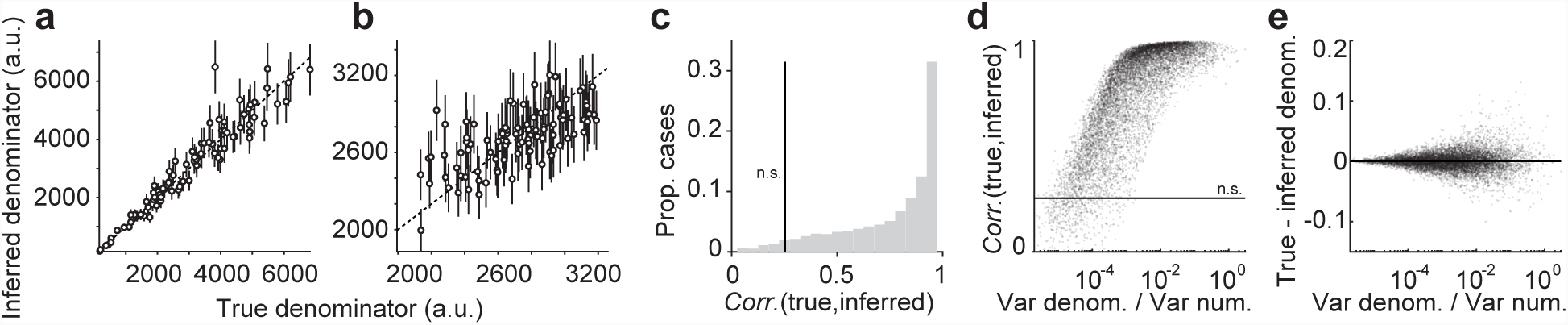
Inference of single-trial normalization strength in the RoG model. (**a**,**b**) Comparison of the true and inferred single-trial normalization strength, across 100 trials of two simulated experiments. Error bars: standard deviation of the inferred values. (**c**) Histogram of Spearman correlation coefficients between true and inferred values, across 10,000 simulated experiments each with 100 trials. The model was formulated as in Methods eq. (1.16) for the contrast response function, with parameters drawn from the uniform distribution in the following intervals: 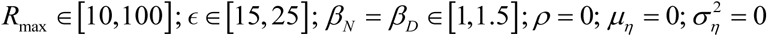, with contrast between 20 and 50%, and with *α_N_*, *α_D_* set to enforce a Fano factor of 1 at a 75% contrast. (**d**) Correlation between true and inferred values (ordinate) as a function of the ratio between the variances of the denominator and numerator in the RoG model (abscissa). Correlation coefficients smaller than the vertical line (**c**) or horizontal line (**d**) are not statistically significant (p>0.05). (**e**) Difference between true and inferred values of the denominator, expressed as a fraction of the true value.

This analysis suggests that for neurons and stimulus conditions with a relatively large variance of *D*, we can infer accurately the single-trial value of *D*. Out of the 152/172 neurons for which the RoG fit quality was larger than 0.5, we found that n=81 neurons satisfied this criterion (specifically, that the variance of *D* was at least 10% of the variance of *N*) for at least one stimulus contrast level. For those cases, we predicted that, if the inference is accurate, during trials with larger normalization signal the response variability should be lower. To test this prediction, we estimated *D* in each trial, and then split trials within each contrast condition in two sets containing values of *D* either above or below average (note that, in many cases, the set of trials with *D* below average contained no spikes at all; therefore, to obtain meaningful estimates of Fano factors, we excluded those cases, leaving us with n=39 neurons). As a control, we verified that in the trial set with large inferred *D*, the mean spike count was always lower (Fig. 5a), indicating that our estimates of *D* were meaningful. Importantly, both the spike count variance and the Fano factor were also systematically lower for the large-*D* trials (Fig. 5b,c), in agreement with the prediction of the RoG model. Across neurons and conditions, we observed a 44% reduction (c.i. [33 53]%) in Fano factor for large-*D* versus small-*D* trials. This result was robust to the choice of threshold, i.e. the minimum ratio between the variances of *D* and *N* (Fig. 6a-c). The changes were absent if we randomly split trials in two groups (average difference in Fano over 1,000 splits = −2%, c.i. [-8 3]%). As a further control, we verified that if we instead considered only cases in which the ratio between the variances of *D* and *N* was less than a small threshold (i.e. where we expect poor single-trial estimates of *D*), there were no significant differences in Fano factor between large-*D* and small-*D* trials, unless we made the threshold large enough to effectively include many cases with relatively large variance of *D* (Fig. 6d-f).

**Figure 5.**
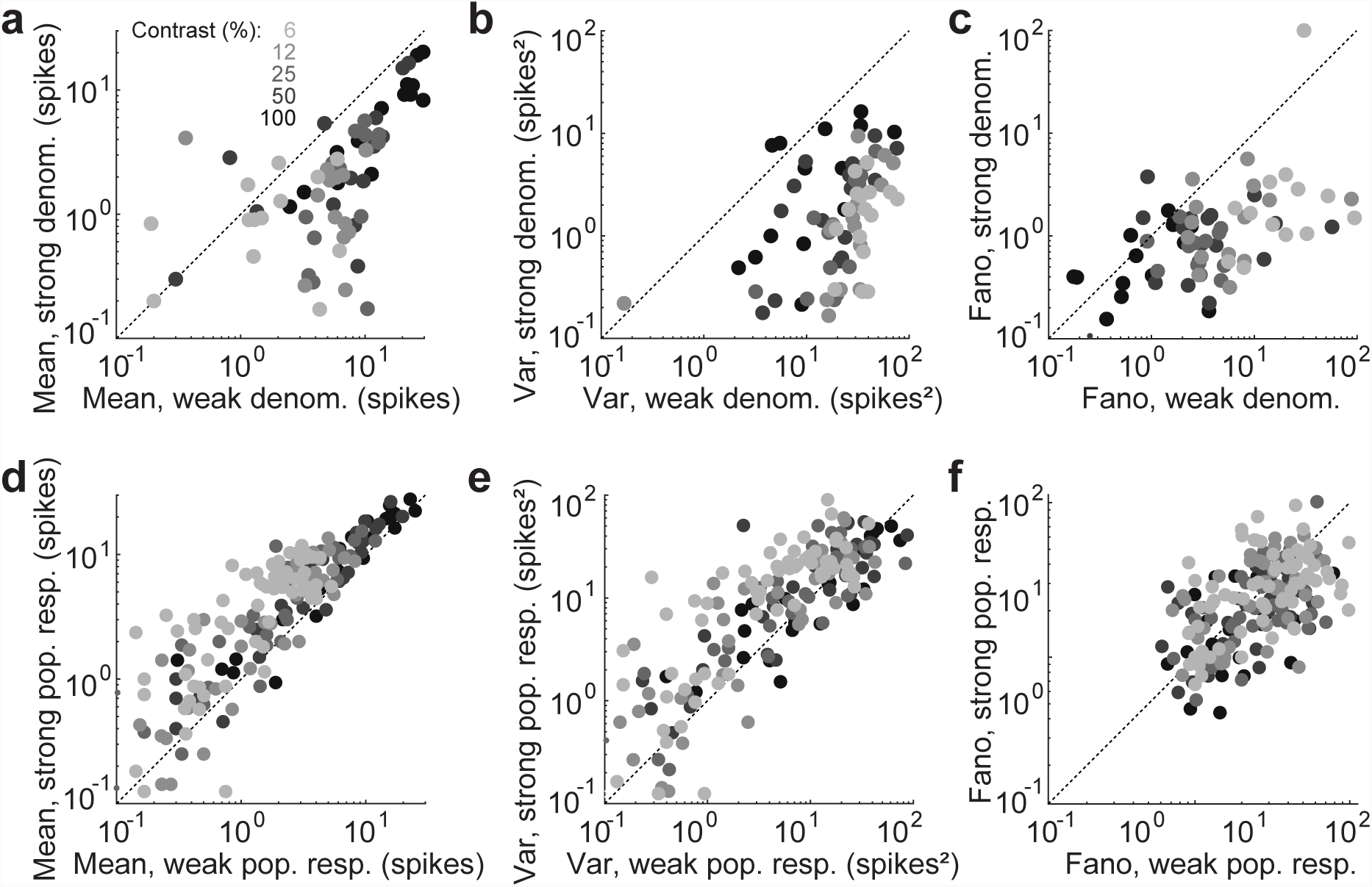
Reduced response variability during epochs with strong normalization. (**a**) Each symbol denotes, for one neuron and one contrast condition, the mean spike count across trials with inferred strong (ordinate) versus trials with inferred weak (abscissa) normalization signal. Only neurons with large across-trials variance of the normalization signal (at least 10% of the variance of the numerator), and contrast conditions with at least one spike in each subset of trials, are included. (**b,c**) Same as (**a**) but for response variance (**b**) and Fano factor (**c**). (**d**,**e**,**f**) Same as (**a,b,c**) but using as a proxy of single-trial normalization strength the total activity of a population of simultaneously recorded neurons.

**Figure 6.**
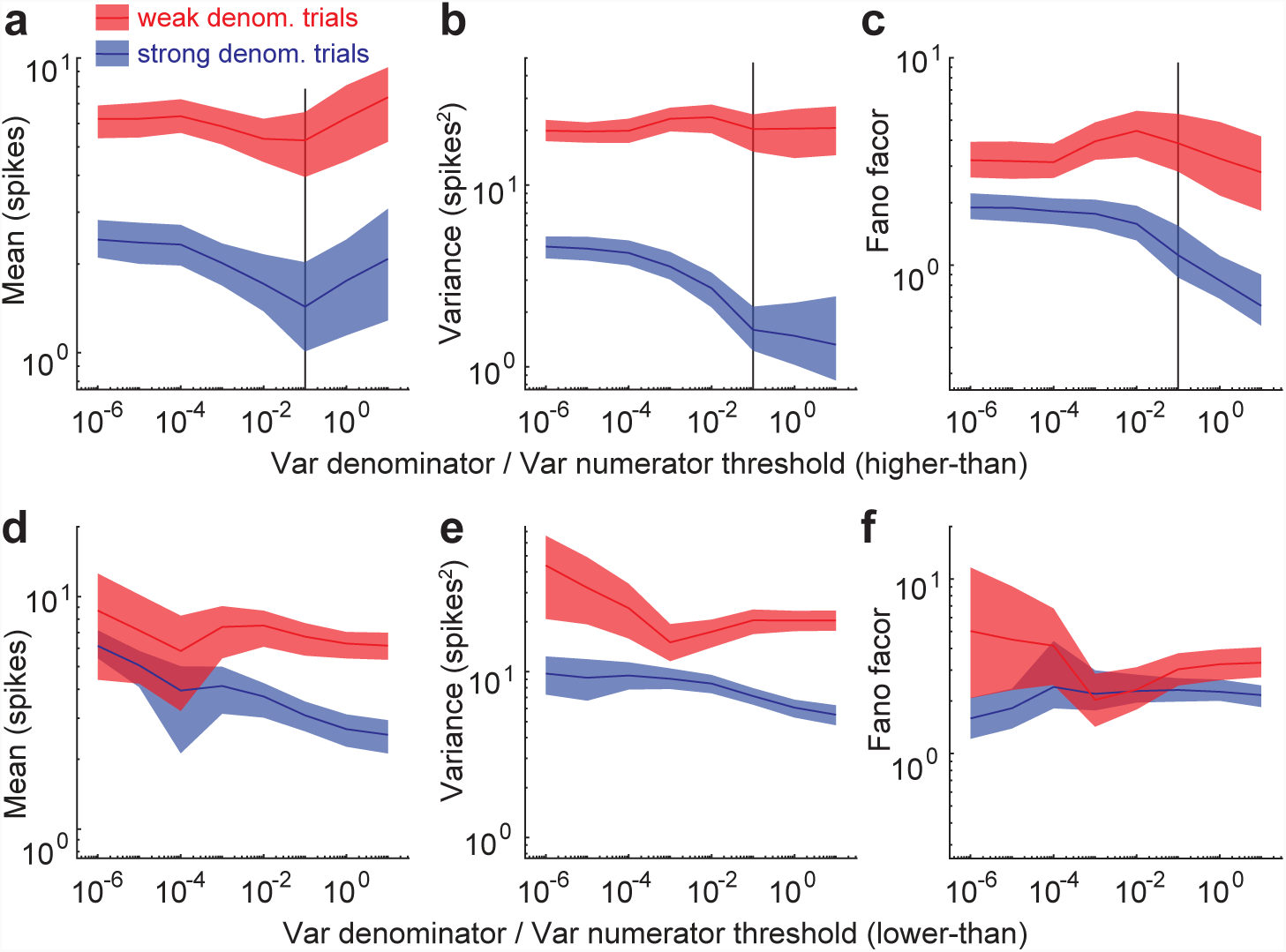
Reduced variability during epochs with strong inferred normalization is robust to choice of threshold. (**a**) Average spike count (ordinate) across neurons and contrast conditions for which the ratio between denominator variance and numerator variance was larger than the threshold (abscissa). Averages are computed over the subset of trials with strong (blue) or weak (red) inferred normalization signal. Shaded areas: 95% c.i. Black vertical line: the threshold used for Fig. 5. (**b,c**) Same as (**a**) but for the spike count variance (**b**) and Fano factor (**c**). (**d-f**) Same as (**a-c**), but including neurons and contrast conditions for which the ratio between denominator variance and numerator variance was smaller than the threshold (abscissa). The largest threshold in (**d-f**) and the smallest threshold in (**a-c**) correspond to including all neurons and contrast conditions.

We next asked whether the total population activity may be a good proxy for the single-trial normalization strength, as normalization is thought to result from pooled activity of many neurons (Carandini & Heeger, 2011; Heeger, 1992). However, when we split trials based on total population activity we found larger mean spike count for trials with larger population activity (an average increase of 28%, c.i. [24 31]%; opposite to what would be predicted by stronger normalization) and minimal change in Fano factor (a 5% reduction, c.i. [1 9]%; Fig. 5d-f). This may reflect that, whereas the normalization strength is conceptualized as the total activity of the un-normalized population, the measured population activity is already normalized and it is therefore not a good measure of normalization strength. Alternatively, it may be that the normalization pool is distinct, or much larger, than the neurons we could record from simultaneously.

These results thus demonstrate that the RoG model enables accurate inference of the single-trial normalization strength for a subset of neurons. Using this method, we found that response variability was lowered during epochs with strong normalization.

## Discussion

Divisive normalization is ubiquitous in neural computation, and can dramatically affect both the strength and the variability of neural activity. Existing models do not offer a way to quantify the influence of normalization on variability. Here we have introduced a simple yet powerful model, the RoG, that extends the standard normalization model to variable neuronal activity. The RoG offers two major advantages over existing models of variability: first, the explicit normalization formulation allowed us to study directly the impact of normalization on response variability; second, the RoG treats the normalization signal as a hidden random variable, which allowed us to estimate the single-trial normalization strength via Bayesian inference.

We have shown that the RoG describes accurately the contrast dependence of spike count mean and variance in macaque V1. In particular, the model was able to reproduce the systematic reduction of the Fano factor as contrast increased, which we observed in the majority of recorded neurons (Fig. 2c,d). This effect is consistent with similar recent observations (Orbán et al., 2016), but it could not be captured by the modulated-Poisson model (Fig. 2e), a popular framework to describe response variability (Goris et al., 2014). Recently, another model has been introduced that extends the classical Linear-Nonlinear-Poisson (LNP) framework, termed flexible overdispersion (Charles et al., 2018), that can capture a decrease of Fano factor as firing rate increases, and thus could fit well our observation on contrast response functions. However, this model uses a simple form of response nonlinearity as is common in the LNP framework to make the optimization tractable, and so it would not allow to study the role of divisive normalization. One possibility would be to treat both the numerator and denominator of the standard normalization model as LNPs, i.e. similar to the RoG but using LNPs instead of Gaussians. The disadvantage of this approach is that the mathematical result of (Marsaglia, 1965) would not apply: the spike count distribution may not be well approximated by a Gaussian, and one would no longer have simple expressions for the spike count mean and variance as in eq. (1.4), nor closed-form, instantaneous inference of the single-trial normalization signal as in eq. (1.12).

Thanks to the explicit normalization in the RoG, analysis of the model indicated a general link between normalization and variability (Fig. 3a). Consistent with this, we observed that neurons with stronger normalization were relatively less variable (Fig. 3b-d), and also that for most neurons variability was reduced during response epochs in which normalization was stronger than average (Fig. 5a-c). One possible interpretation of these observations is that normalization may reflect a general mechanism to stabilize cortical activity, for instance one based on dynamic balance between excitation and inhibition (Adesnik, 2017; Hennequin et al., 2018). However, diverse influences of normalization on variability may be apparent across cortical areas or with different experimental manipulations, possibly reflecting the diverse network mechanisms thought to underlie the effects described by normalization (Carandini & Heeger, 2011), including feedforward (Carandini et al. 2002; Webb et al. 2005), recurrent (Ozeki et al. 2009; Rubin et al., 2015) and feedback (Nassi et al. 2013; Nurminen et al. 2018) signals. Although our analysis revealed primarily a stabilizing effect of normalization, the RoG model is flexible enough to capture also the opposite effect.

Two recent studies have also pointed to a possible impact of normalization on response variability (Tripp, 2012; Verhoef & Maunsell, 2017). Those studies focused on variability shared between neurons, i.e. noise correlations, and relied exclusively on simulations. In contrast, the RoG framework allowed us to derive analytical results linking normalization and variability. Furthermore, the explicit probabilistic formulation and simple parametric form of the RoG allows for quantitative data fitting and single-trial inference, which are not tractable in the models of (Tripp, 2012; Verhoef & Maunsell, 2017). The RoG model could also be extended to capture the joint responses of two or more neurons, simply by including *N* and *D* terms for each neuron, and assuming a multivariate Gaussian distribution for the resulting collection of variables (i.e. with dimension 2 times the number of neurons). With this extension, the number of parameters (i.e. the covariance matrix of the multivariate Gaussian) would grow quadratically with the number of neurons, thus potentially limiting the applicability or requiring strong regularization for large numbers of neurons. However, for pairwise responses and under the assumption of independence between *N* and *D* (which was supported by our data, see Methods), the pairwise RoG would only require two additional parameters for the correlation between the *N* terms and between the *D* terms.

To optimize the RoG parameters on data, it is necessary to consider stimulus manipulations for which the firing rate is well described by the standard normalization model. This is the case for contrast response functions, which motivated our choice here to validate the RoG model. There are other stimulus manipulations that are thought to involve normalization in V1, as observed in phenomena like cross-orientation masking (Bonds, 1989; Carandini et al., 1997; DeAngelis et al. 1992; Schwartz & Simoncelli, 2001), surround suppression (Cavanaugh et al. 2002; Coen-Cagli et al., 2015; Schwartz & Simoncelli, 2001), and temporal adaptation (Heeger, 1992; Snow et al. 2016; Solomon & Kohn, 2014; Westrick et al. 2016). Furthermore, normalization is also widespread (Carandini & Heeger, 2011) in other visual areas (Rust et al. 2006), and in other sensory (Ohshiro et al. 2011; Olsen et al. 2010; Rabinowitz et al. 2011) and non-sensory (Louie et al. 2014) areas. Applying the RoG model to those datasets could therefore reveal whether normalization has an impact on variability in other domains, similar to what we have found in V1 for contrast manipulations.

Note that the logic of the model proposed here is fundamentally different from that of models proposed by us and others to explain the widespread presence of normalization (Beck et al. 2011; Coen-Cagli et al. 2012; Coen-Cagli et al., 2015; Lochmann et al. 2012; Schwartz et al. 2007; Schwartz & Simoncelli, 2001; Sinz & Bethge, 2013; Snow et al., 2016; Wainwright et al. 2001). Models in this class hypothesize a specific goal of neurons (e.g. efficient coding of sensory inputs (Barlow, 1961)) and ask whether normalization achieves that goal, i.e. these models aim to explain why neurons behave in a way that is well described by normalization. Our goal here instead was to provide a descriptive model, i.e. a model that parametrizes the effects of normalization on neural responses in a way that is easy to fit to data, allowing us to quantify and interpret those effects. Using this approach, we have revealed an apparent relation between normalization and variability; explaining why this relation exists and what functional role it may play, falls in the domain of the other class of models (e.g. (Orbán et al., 2016)).

Although we focused on contrast modulation, another case of particular interest is temporal adaptation, as it has been hypothesized that adaptation effects reflect a modulation of the normalization signal itself (Aschner et al. 2018; Heeger, 1992; Solomon & Kohn, 2014; Wainwright et al., 2001; Westrick et al., 2016). Inference of the single-trial normalization signal, as enabled by the RoG model, would allow testing this hypothesis directly. However, one limitation of the single-trial inference of normalization strength is that it is only reliable for neurons that have a relatively large across-trial variance of the normalization signal. In our dataset, only about half the neurons satisfied this criterion (81/152). Such a small percentage may reflect the fact that the effective normalization signal results from the pooled activity of many neurons, as commonly assumed in the normalization framework (Carandini & Heeger, 2011). If neurons in the normalization pool have homogeneous response properties, pooling their responses should produce a more stable signal than the excitatory drive to one neuron. This explanation also suggests that neurons with a relatively larger variance of *D*, according to the RoG, may be normalized by a more heterogeneous normalization pool. This could be tested by measuring the tuning of the normalization signals (Webb et al., 2005) and studying if normalization tuning width correlates with the estimated variance of the normalization signal. Furthermore, this explanation suggests that it should be possible to differentially adapt the normalization pools of the two classes of neurons: for instance, a narrow-band adapter would weaken normalization effects more for the neurons with small variance of *D* according to the RoG.

Single-trial inference of normalization could also prove useful in understanding attentional modulation of perception. It has been suggested that attention improves perceptual decision-making by modulating correlated response variability (Cohen & Maunsell, 2009; Mitchell et al., 2009), and that this modulation results from a dynamic mechanism for normalization (Verhoef & Maunsell, 2017). Our method could be extended to pairwise responses and noise correlations, and would then allow one to test this hypothesis directly by relating single-trial normalization strength and behavioral performance.

## Materials and Methods

### The Model

#### The Divisive Normalization model and the RoG model

In the generic formulation of the normalization model, the average spike count of a neuron is described by:

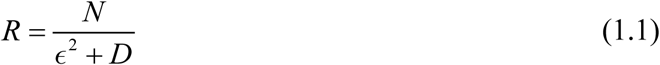

where the numerator *N* usually represents the stimulus drive to the neuron; *D* is the normalization signal, typically conceptualized as the summed activity of a large group of neurons termed normalization pool; and the constant ϵ prevents division by zero.

We extended the normalization model to account for response variability by treating both *N* and *D* as Gaussian-distributed random variables (Fig. 1a), and adding Gaussian noise to the ratio (to capture spontaneous activity fluctuations), leading to the RoG model:

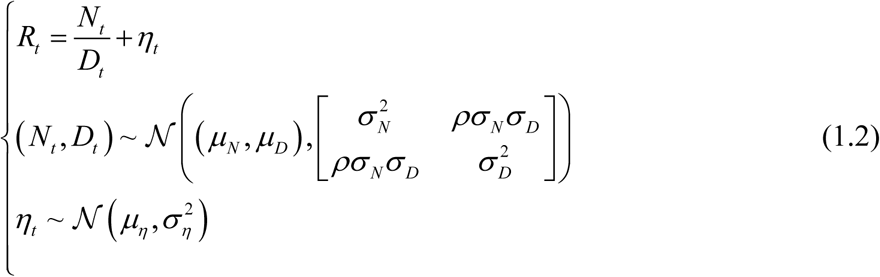

where the subscript *t* indexes a single trial. Note that we have implicitly absorbed _ω_^2^ in *μ*.

#### Approximated moments of the RoG distribution

Analytical expressions for the RoG distribution (Hinkley, 1969; Pham-Gia et al., 2006) and other ratio distributions (Pham-Gia & Turkkan, 2005; Springer, 1979) have been obtained, but they involve complicated expressions that make it difficult to perform optimization and inference, and to find the moments of the distribution. A well-known example is the ratio of two standard normal variables, i.e the Cauchy distribution, whose mean and variance are not even defined. This is important because we are interested in characterizing the effects of normalization on response variability. For our purposes, a useful result is that, if *D* has negligible probability mass at or below 0, the RoG is closely approximated by a Gaussian (Marsaglia, 1965, 2006; Pham-Gia et al., 2006). In our context, this assumption on *D* is well motivated, as *D* is usually thought of as the summed activity of a large normalization pool of neurons, and so unlikely to be zero or negative. Under these conditions, the RoG output is Gaussian distributed and therefore fully characterized by its mean and variance, for which we can derive approximations using a second order Taylor expansion:

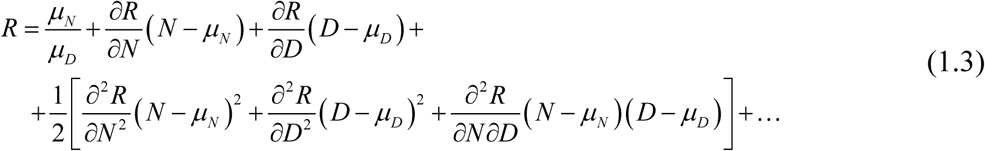

(neglecting higher order terms) and taking the expectations, thus obtaining the following expressions:

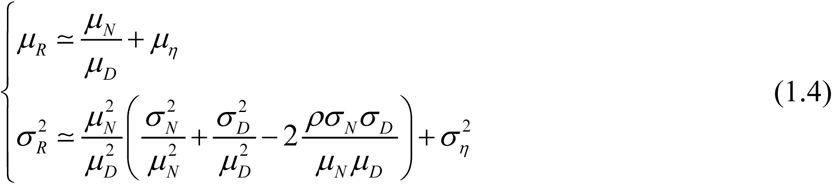

where we ignored the second-order term for the mean. We validated the approximations in simulations, and found that both remained unbiased (within 0.3%) over several orders of magnitude (Fig. 1b).

#### Analytical relation between spike-count variance, stimulus drive, and normalization strength

Equation (1.4) allows us to address directly the question of how the average normalization strength (*μ*_*D*_) affects response variability. First, we address the case with no additive noise 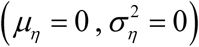. We quantify variability by the Fano factor:

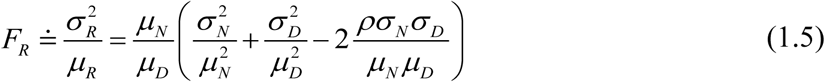

and we study its derivative with respect to *μ*_*D*_:

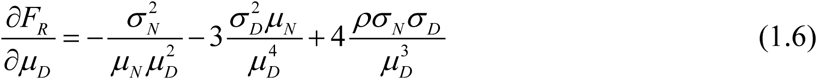

If *N* and *D* are uncorrelated (*ρ* = 0) the derivative is always negative, implying that the Fano factor increases with the strength of normalization. When *ρ* > 0, the derivative can change sign. To find the amount of correlation required for this to happen, we set the derivative to zero:

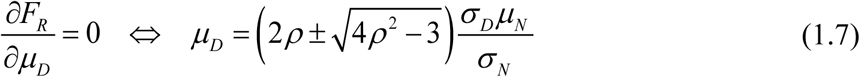

This shows that a solution exists only if *ρ*^2^ ≥ 3/4, therefore an increase in normalization strength can increase the Fano factor only for very large correlation levels between *N* and *D*. Notice also from eq. (1.5) that, regardless of the value of *ρ*, the dependence of the Fano on the numerator mean is not monotonic in general, because it is the sum of a term proportional to *μ*_*N*_ and one proportional to 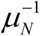.

Next, we consider that case with 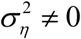. For simplicity we set *μ*_*η*_ = 0, and also *ρ* = 0 because it has a very limited influence per our analysis above. In this case the Fano factor is:

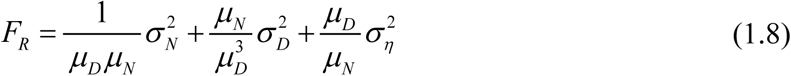

This expression shows that the relative sizes of the additive noise and of the variances of *D* and *N* determine whether the Fano factor increases or decreases as a function of either *μ*_*D*_ or *μ*_*N*_.

#### Reproducing Poisson-like variability

A well-established property of cortical response variability in vivo is that it is Poisson-like, i.e. the variability increases roughly monotonically with the mean spike count (Tolhurst et al., 1983). For the RoG distribution, changes in mean and variance (eq. (1.4)) are controlled by changes in the parameters of eq. (1.2), encompassing a broader range of variance-mean relationships. To further constrain the RoG parameters, we reasoned that both *N* and *D* are described as signals of neural origin, and thus we assumed Poisson-like variability for each of those terms separately, following a commonly used formulation (Carandini et al., 1997; Tolhurst et al., 1983):

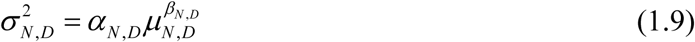

For *β* = 1, the parameter *α* determines the Fano factor, whereas different values of *β* allow for non-constant Fano. Note that we used this relation separately for the numerator and denominator of the RoG model, and this does not imply necessarily that also the variability of *R* follows the same relation. However, we found empirically that for a wide range of parameter settings the variability of *R* is Poisson-like (Fig. 1c). Notice also that, if we substitute eq. (1.9) into eq. (1.5), there is no longer a unique, monotonic relation between the Fano factor and *μ*_*D*_; in particular, for large enough values of *β*_*D*_, the Fano factor can increase (rather than decrease) as *μ*_*D*_ increases.

### Inference of the single-trial normalization signal

An advantage of the Gaussian approximation for the RoG is that it greatly simplifies inference. Specifically, we are interested in inferring the normalization strength *D* in a single trial (which cannot be measured directly), given the measured spike count *R*. To show how this is achieved, here we assume that all the parameters (the means and covariance of *N* and *D*) are known: we explain in the next section the model fitting procedure to estimate those parameters from data.

We are interested in the posterior distribution of *D* given *R*, which can be obtained by Bayes rule:

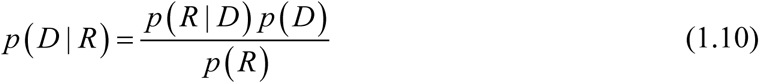

Here *p* (*D*) is the Gaussian prior assumed by the RoG model on *D*. Note that *p* (*R*) does not depend on the value of *D* (only on its mean and variance, which are fixed), therefore we can ignore the denominator. We first consider the case *ρ* = 0 and no additive noise (*η* = 0). Using the fact that 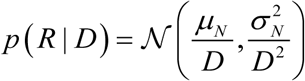, with straightforward calculations we obtain:

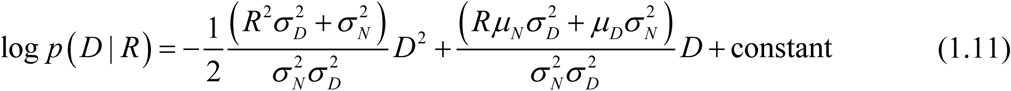

Equation (1.11) is quadratic in *D*, which means the posterior distribution of *D* is Gaussian. It is therefore straightforward to compute the maximum a posteriori (MAP) estimate of *D* which coincides with the posterior mean (i.e. our estimate of the normalization strength, denoted by 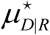), and the posterior variance (i.e. the uncertainty around that estimate, denoted 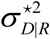):

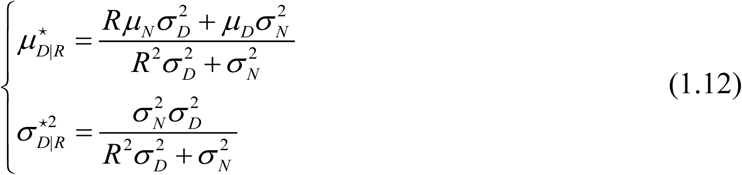

The asymptotic behavior of eq. (1.12) provides some insight into the conditions under which the inference of *D* is informative. In the limit of large prior variance 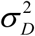 for the denominator (relative to 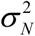), the posterior mean tends to *μ*_*N*_/*R* and the posterior variance to a relatively small value 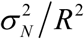. In other words, in this condition, trial-by-trial fluctuations in spike count are entirely explained by fluctuations in the normalization strength, and therefore the normalization strength can be estimated accurately. Conversely, in the limit of large prior variance 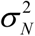 for the numerator, the posterior mean and variance for *D* converge to their prior values, and therefore the estimate of the normalization strength is entirely uninformative (i.e. it is, in each trial, identical to the average across trials).

The case with *ρ* ≠ 0 is similar except that *p* (*R* | *D*), and therefore *p* (*D* | *R*), also depend on *ρ*:

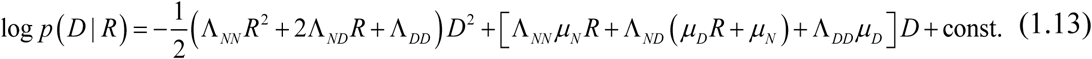

where we denote with Λ the inverse of the covariance matrix between *N* and *D*. As we explain below, in our application to V1 data we found that including *ρ* did not improve the model’s predictive power due to overfitting, therefore we only used eq. (1.12) for the single-trial inference. However using eq. (1.13) would be just as easy in other data that warranted the use of *ρ*.

Unfortunately the case with *η* ≠ 0 does not have such a simple solution. The log-posterior is no longer quadratic in *D*, and to find the MAP estimate one has to set the derivative (with respect to *D*) to zero. We computed the derivative, and found it is a fifth order polynomial, for which the Abel-Ruffini theorem states there is no general solution (Rosen, 1995). The MAP estimate of *D* could instead be found numerically by gradient descent, but we did not pursue this further. Nonetheless, we found in simulations that the estimator of *D* (eq. (1.12)) remains approximately unbiased (within 0.01%) for moderate levels of additive noise, as long as the mean of such additive noise was correctly accounted for (i.e. removing the additive noise mean from the stimulus-evoked spike counts). For this reason, in our analysis of experimental data, we measured the spontaneous rate and removed it from the evoked responses before inferring the single-trial value of *D*. We also verified that our results were noisier, though not qualitatively different, when we did not remove the spontaneous rate.

### Model fitting

#### Maximum likelihood parameter estimation

We optimized the RoG parameters by maximum likelihood. In the general setting, the RoG model has five free parameters (*μ*_*N*_, *μ*_*D*_, *σ* _*N*_, *σ*_*D*_, *ρ*) per stimulus condition, as they can be stimulus dependent, plus (*μ*_*η*_, *σ*_*η*_). The Poisson-like assumption of eq. (1.9) reduces the stimulus-dependent parameters to three (*μ*_*N*_, *μ*_*D*_, *ρ*), plus (*α*, *β*) which are stimulus independent. In our application to the contrast response function data, we found that *ρ* leads to overfitting, therefore we set it to zero, and we further parametrized the stimulus-dependence of the means (details below). If we denote generically the free parameters by Φ, the log-likelihood of a dataset of measured responses {*R*_*t*_ (*s*)} (where *t* denotes the trial and *s* the stimulus value) is

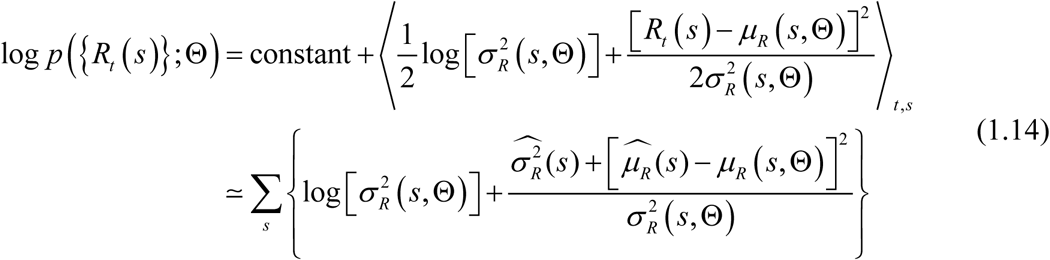

where in the second line we have dropped the constant; ⟨. ⟩_*t,s*_ denotes the expectation over trials and stimuli; 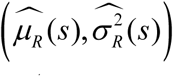denote the empirical mean and variance across trials, per stimulus condition; and 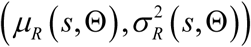 are the model predictions. To find the best fit parameter values, we minimize the negative of the log-likelihood iteratively using Matlab’s ‘fmincon’.

By inspecting eq. (1.4) for the variance, it can be noted that the additive noise acts as a regularizer in eq. (1.14), i.e. preventing the model variance to become too small. One important difference between fitting the RoG and fitting the standard normalization model, is that in the RoG both the predicted mean and the variance of the spike count are stimulus-dependent and so both appear in the objective function, whereas in the standard model only the mean is stimulus-dependent and the optimization reduces to a nonlinear least-squares problem.

#### Cross-validated goodness of fit

We quantified the ability of the model to capture the data on a test dataset distinct from the training set used for optimization. The training set contains all stimulus conditions and all but one repetition; this held-out repetition, for all stimuli, forms the test set. We performed leave-one-out cross validation, i.e. we averaged the test-set results across all possible splits of the data between training and test set. The measure of goodness of fit (a.k.a. predictive power) is the average log-likelihood computed on the test data. To map this value on an interpretable scale, we normalized it between a null model and an oracle model as is done often (Goris et al., 2014; Stocker & Simoncelli, 2006). The null model predicts that the response mean and variance are stimulus independent and equal to the mean and variance across all stimuli and repetitions in the training set. The oracle model predicts that the mean and variance in each stimulus condition are identical to the means and variances estimated from the training set. The normalized, cross-validated goodness of fit approaches zero when the model is as bad as the null model, and approaches 1 when it is as good as the oracle.

#### Parametrization of the contrast response function

To fit the RoG model, we need to specify how 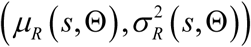depend on the stimulus and the parameters, which in turn requires parametrizing the stimulus-dependence of the moments of *N* and *D*. Here we apply the RoG to contrast response function data. Therefore, we start from the standard normalization formulation which provides excellent fits to the average spike count as a function of contrast (Heeger, 1992):

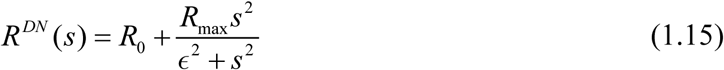

Here, *s* denotes contrast (assuming the orientation of the visual stimulus is fixed), and (*R*_0_, *R*_max, ω_) are free parameters. Based on eq. (1.15), we parametrize the RoG model such that the predicted mean response (eq. (1.4)) is identical to the prediction of the normalization model:

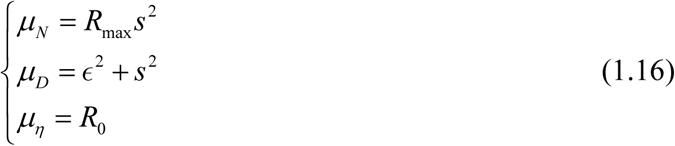

and we have additional free parameters for the noise variance 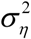, as well as (*ρ,α* _*N*_,*β*_*N*_,*α*_*D*_,*β*_*D*_ from eq. (1.2) and eq. (1.9). We set *R*_0_ to the average spontaneous activity, and we constrain the parameter range as follows: ∫ ∈[1,100]; *α*_*N, D*_ ∈[.1, 20]; *β*_*N, D*_ ∈[1, 2]; *ρ* ∈[-1,1]; *R*_max_ between 0.5 and 2 times the maximum response across all stimuli; and 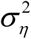 between 0.1 and 10 times the measured variance of the spontaneous activity. In practice, we found that treating 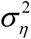 as a free parameter slightly improved predictive power while changing little from its measured values. On the contrary, treating *ρ* as a free parameter produced strong overfitting (average goodness of fit on the training set 0.82, c.i. [0.8 0.85]; on the test set 0.65, c.i. [0.64 0.68]). Therefore, we report below the results with *ρ* = 0 rather than as a free parameter.

### Alternative Models

The model comparison of Fig. 2 uses the popular Modulated-Poisson model (Goris et al., 2014), which has been shown to capture well the deviations of cortical variability from Poisson statistics (i.e. from constant Fano factor of 1). In brief, the Modulated-Poisson model assumes that spike counts evoked by a fixed stimulus are Poisson distributed, with a rate determined by the product of a constant stimulus function (the tuning curve) and a fluctuating gain factor (termed modulator). Across-trials fluctuations of the modulator induce super-Poisson variability, leading to Fano factors that increase with the mean spike count. In our Fig. 2, we used the model of (Goris et al., 2014) for the spike count distribution, but instead of the measured tuning curves we used the divisive normalization model of eq. (1.15) as the tuning curve. We then fit jointly all parameters (those of the normalization model for the mean tuning curve, plus the variance of the gain factor). Note that this formulation of the Modulated-Poisson model has fewer free parameters than the RoG. For this reason, we compare the two models by cross-validation, which automatically accounts for the difference in model complexity.

### Experimental Data and Statistical Analysis

#### Animal preparation and data collection

Data were collected from four adult male monkeys (Macaca fascicularis). Animal preparation and general methods were described previously (Jia et al. 2013). In brief, anesthesia was induced with ketamine (10 mg per kg of body weight) and maintained during surgery with isoflurane (1.0–2.5% in 95% O2). During recordings, anesthesia was maintained by sufentanil citrate (6–18 μg per kg per h, adjusted as needed for each animal). Vecuronium bromide (0.15 mg per kg per h) was used to minimize eye movements. All procedures were approved by the Albert Einstein College of Medicine at Yeshiva University and followed the guidelines in the United States Public Health Service Guide for the Care and Use of Laboratory Animals.

We recorded neuronal activity using arrays of 10 × 10 microelectrodes (400-μm spacing, 1-mm length) inserted in the opercular region of V1. Waveform segments that exceeded a threshold (a multiple of the RMS noise on each channel) were digitized (30 kHz) and sorted off-line. For all analysis we included signals from well-isolated single units as well as small multi-unit clusters, and refer to both as neurons.

#### Visual stimuli and presentation

We displayed stimuli on a calibrated CRT monitor (1,024 × 768 pixels, 100-Hz frame rate, ∼40 cd m^-2^ mean luminance) placed 110 cm from the animal, using custom software. We first measured the spatial RF of each neuron, using small gratings (0.5 degrees in diameter, four orientations, 250-ms presentation) presented at a range of positions. Stimuli for the main experiment were then centered on the aggregate RF of the population.

To measure contrast response functions, we presented drifting gratings (1-degree diameter) at 4 orientations and 5 contrast levels (6.25, 12, 25, 50, 100)%. We also added a 0-contrast condition to measure spontaneous activity. Stimuli were presented for 500 ms (3 animals) or 800 ms (1 animal), separated by a blank screen of equal duration, repeated 20-25 times and interleaved in pseudo-random order. For each neuron, we analyzed contrast response functions only at the best orientation tested.

#### Characterization of neuronal responses and inclusion criteria

We quantified neuronal activity on each trial by the spike count measured in a window starting 50 ms following stimulus onset (to account for response latency) and ending 50 ms after stimulus offset. Due to the small stimulus size (1 degree) relative to the typical spread of RF centers on the array (∼2.5 degrees at the recorded eccentricity), many neurons were not driven by the stimuli. Therefore, we included in the analysis only neurons that were visually responsive (n=172), defined by their average firing rate to the best stimulus being larger than the spontaneous firing rate plus two standard deviations.

#### Statistical analysis

Confidence intervals reported throughout the text are 95% intervals estimated on 10,000 bootstrap samples. When averaging Fano factor across neurons or conditions, we computed the geometric mean. Significance and p-values were obtained with parametric t-test when comparing goodness-of-fit for different models and in Table 1, and with a permutation test for the Spearman correlation coefficient.

## Acknowledgements

We thank A. Kohn for helpful discussions and for comments on the manuscript. S.S.S. gratefully acknowledges financial support from Fight for Sight. The data were collected in the Kohn lab, with support from NIH grants (EY026240 and EY016774).

